# Mathematics and the brain –a category theoretic approach to go beyond the neural correlates of consciousness

**DOI:** 10.1101/674242

**Authors:** Georg Northoff, Naotsugu Tsuchiya, Hayato Saigo

## Abstract

Consciousness is a central issue in cognitive neuroscience. To explain the relationship between consciousness and its neural correlates, various theories have been proposed. We still lack a formal framework that can address the nature of the relationship between consciousness and its physical substrates though. Here, we provide a novel mathematical framework of Category Theory (CT), in which we can define and study the “sameness” between “different” domains of phenomena such as consciousness and its neural substrates. CT was designed and developed to deal with the “relationships” between various domains of phenomena. We introduce three concepts of CT including (i) category; (ii) inclusion functor and expansion functor; and (iii) natural transformation between the functors. Each of these mathematical concepts is related to specific features in the neural correlates of consciousness (NCC). In this novel framework, we will examine two of the major theories of consciousness: integrated information theory (IIT) of consciousness and temporo-spatial theory of consciousness (TTC). These theories concern the structural relationships among structures of physical substrates and subjective experiences. The three CT-based concepts, introduced in this paper, unravel some basic issues in our search for the NCC; while addressing the same questions, we show that IIT and TTC provide different albeit complementary answers. Importantly, our account suggests that we need to go beyond a traditional concept of NCC including both content-specific and full NCC. We need to shift our focus from the relationship between “one” neuronal and “one” phenomenal state to the relationship between a structure of neural states and a structure of phenomenal states. We conclude that CT unravels and highlights basic questions about the NCC in general which needs to be met and addressed by any future neuroscientific theory of consciousness.

**Author summary:** Neuroscience has made considerable progress in uncovering the neural correlates of consciousness (NCC). At the same time, recent studies demonstrated the complexity of the neuronal mechanisms underlying consciousness. To make further progress in the neuroscience of consciousness, we need proper mathematical formalization of the neuronal mechanisms potentially underlying consciousness. Providing a first tentative attempt, our paper addresses both by (i) pointing out the specific problems of and proposing a new approach to go beyond the traditional approach of the neural correlates of consciousness, and (ii) by recruiting a recently popular mathematical formalization, category theory (CT). With CT, we provide mathematical formalization of the broader neural correlates of consciousness by its application to two of the major theories, integrated information theory (IIT) and temporo-spatial theory of consciousness (TTC). Together, our CT-based mathematical formalization of the neural correlates of consciousness including its specification in the terms of IIT and TTC allows to go beyond the current concept of NCC in both mathematical and neural terms.

## Introduction

### “There is no certainty in sciences where mathematics cannot be applied” (Leonardo da Vinci)

Consciousness has long been regarded as a mysterious phenomenon, and it was mainly dealt within philosophy. Past philosophers like Descartes argued that consciousness is only accessible from the first-person perspective and cannot be explained from the third-person perspective. This tradition is followed by present philosophers who speak of an unbridgeable gap between the third-person physical objects of brain and first-person consciousness – formulated as the “explanatory gap problem” [64] or the “Hard problem” [17] (see Part IV in Northoff 2014b for a general overview). However, recent neuroscientific research demonstrates that consciousness is a biological phenomenon and the first-person perspective and phenomenal experience needs to be explained in the scientific framework [19, 72, 73, 75, 94].

The recognition of consciousness as a biological phenomenon has led neuroscience to search for the neural correlates of consciousness (NCC; [4, 21, 22, 59, 71, 72]). The NCC has been defined as the minimum neuronal mechanisms jointly sufficient for any one specific conscious percept [59]. Recent progress in consciousness research further introduces two refined interpretations of the NCC as 1) content-specific NCC, which determines a particular phenomenal distinction with an experience and 2) full NCC, which supports conscious experiences in their entirety, irrespective of the contents [60][1].

Major neuroscientific theories of consciousness, based on the empirical neuroscientific findings around the NCC, include the Integrated Information Theory (IIT) [101, 102], the Global Neuronal Workspace Theory (GNWT) [23-25] and most recently, the Temporo-spatial Theory of Consciousness (TTC) [70, 72, 73, 77]. Others include the higher order theories of consciousness [63,87], recurrent processing theory [62], operational space and time [33], neural synchrony [30] and social and attention schema theory [37]. As the discussion of all these approaches is beyond the scope of this paper, we focus on two of the major theories, the Integrated Information Theory (IIT) and Temporo-spatial Theory of Consciousness (TTC).

The essential problem in our search for the NCC consists in bridging two domains of ‘relationships’: relationships among the contents in conscious experience in the mental domain and relationships among neurons in the physical domain. One can thus speak of “neuro-phenomenal relationship” connecting the brain’s neuronal states and the phenomenal features of consciousness [72, 75]. Or, presupposing a broader more ontological context, one can speak of an “identity between experiences and conceptual structures” [102, p.11]. Independent of how one frames the relationship in conceptual terms, theories about the NCC must address this fundamental problem about the ‘relationships’ between physical and mental domains [105]. Transcending the empirical investigation of the neuronal states themselves, this requires mathematical tools to formalize the ‘relationships’ between the two domains. The objective of our paper is to provide a first step towards developing a mathematical formalization of the relationship between neuronal and phenomenal domains in the NCC. This will first be explicated on mathematical grounds and then applied to the NCC with IIT and TTC serving as paradigmatic test cases.

In consciousness research, there has been sporadic attempts to apply mathematical tools to bridge the gap between the physical and the mental domains (Just to name a few, [Stanley 1999 J of consciousness studies, p49-60, Yoshimi 2011 Frontiers in Psychology, Hoffman 1980 Mathematical Modeling pp349-367, Hoffman 1966 J of Mathematical Psychology 65-98, Palmer 1999 BBS]). However, tools such as graph theory, topology, algebra and set theory are not sufficient to deal with the problem of consciousness. What is lacking in these mathematical tools is strong mathematical formalization of “relationships”. Because the relationships are so fundamental in both in the physical and the mental domains, the mathematical tools that are built to deal with the “relationships” is the ideal tool for the studies of the NCC. Here, we introduce another mathematical formalism, called Category Theory (CT). CT provides us with rich and mathematically well-developed classes of relationships.

Historically, CT was developed to establish and formalize relationships between different domains of knowledge that seem to differ in a fundamental way (for example, the mathematical field of algebra and geometry) [29]. Such relationship could be established by introducing the notion of “natural equivalence”. Recently, CT has been proven extremely successful in connecting distinct domains of knowledge as when unifying quantum mechanisms, topology, logics, and computation [7]. That renders CT a suitable mathematical candidate for consciousness research in its quest to formalize the relationship between two distinct domains, the physical and the phenomenal.

In fact, CT has been applied in neuroscience to memory [26-28] neural networks [44], perception [4], cognition [82, 83]. Going beyond a previous more general first attempt [105], we here propose that CT provides a useful mathematical framework for formalizing the “neuro-phenomenal” [72, 75] relationship that underlies consciousness. For that purpose, we introduce three core concepts of CT including (i) category; (ii) inclusion functor and expansion functor; and (iii) natural transformation between them.

We introduce these mathematical concepts to help formalize and account for the neuro-phenomenal relationship. This will be explicated in the context of TTC and IIT. We conclude that these CT-based concepts highlight similarities and complementarities in IIT and TTC. Importantly, CT unravels and highlights several conceptual problems about both content-specific and full NCC. In short, we point out that exclusive focus on the relationship between one neuronal and one phenomenal state is unlikely to yield further fundamental progress in neuroscience of consciousness. Rather, we suggest that the focus should be on the relationships between different neuronal states and different phenomenal states. Such a shift of the focus will naturally lead to future neuroscientific theories of consciousness, which extend and go beyond the traditional concept of the NCC.

## Category and consciousness

### Definition of category

A category is a system consisting of objects and arrows and satisfying the following four conditions (Fig 1).

**Fig 1.**
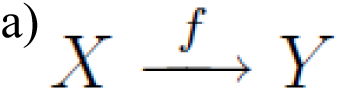
a) Objects, arrows, domain, codomain. Each arrow f is associated with two objects dom(f) and cod(f), which are called domain and codomain of f. When dom(f) = X and cod(f) = Y, we denote f:X →Y, as shown in Fig 1a. (The direction of the arrow can be in any direction, from left to right or reverse, whichever convenient.) A system with arrows and objects are called diagrams.

**Fig 1.**
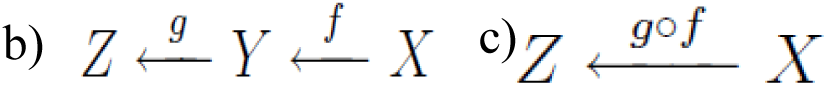
b) Composition. If there are two arrows f and g, such that cod(f) = dom(g), in other words, there is a unique arrow, **c)** g∘f, called the composition of f and g. A diagram is called commutative when any compositions of arrows having the common codomain and domain are equal.

**Fig 1.**
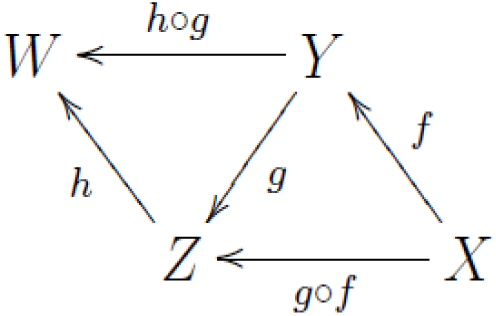
d) Associative law. (h∘g)∘f=h∘(g∘f). In other words, the diagram is commutative.

**Fig 1.**
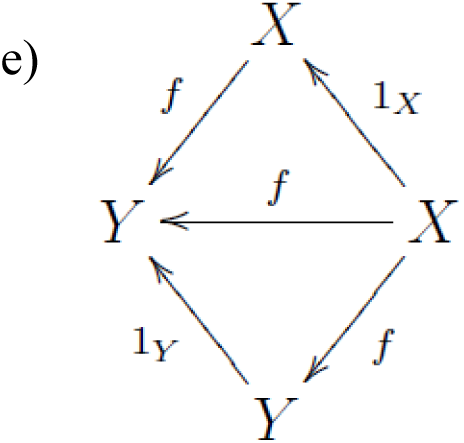
e) Unit law. For any object X there exist an arrow 1x:X→X. Such that the diagram is commutative for any f: X → Y. In other words, f∘1X = f = 1Y∘f for any f .1x is called the identity of X.

By the natural correspondence from objects to their identities, we may “identify” an object (e.g., X) as its identity (e.g., 1x). In other words, we may consider objects are just special cases of arrows. This is one exemplar case where arrows play a more important role than objects in CT. In the following we sometimes adopt this viewpoint.

To sum up, the formal definition of a category is as follows:

#### Definition 1

*A category is a system composed of two kinds of entities called objects and arrows, which are interrelated through the notion of domain/codomain, equipped with composition and identity, satisfying the associative and the unit law.*

One of the strengths of the category theory is that it provides a unified formulation of sameness between different things, based on the notion of isomorphism, which is “invertible” arrows. More precisely:

#### Definition 2

*An arrow f: X* → *Y in a category C is called an isomorphism in C if there exist some arrow g: Y* → *X such that g∘f = 1X and f∘g = 1Y. Two objects are called isomorphic if there is some isomorphism between them.*

Two isomorphic objects are “essentially the same” within the category. If X and Y are isomorphic and X are linked to some other objects through some arrow, then composition with the isomorphism provides the arrow from Y as well. Then Y can be considered as version of X, which is “the same as” X, even when Y is completely different from X.

A famous isomorphism is “the sameness” between a donut and a coffee cup in topology. It actually means that they are isomorphic in a category Top, whose objects are topological spaces (a vast generalization of the notion of “figures”) and arrows are continuous map, i.e., continuous transformation. We will use this notion isomorphism in the following sections.

### Category and consciousness

One of the most fundamental problems in consciousness research is to clarify the “relationship” between the neuronal and the phenomenal domains, the neuro-phenomenal relationship as stated by TTC [72, 77].

From the CT viewpoint, the phenomenal domain can be formulated as a category whose objects are contents of consciousness as experienced and arrows are relationships between them as experienced. Let us call this category “the phenomenal category” and denote it as P.

The formulation of “the neuronal category” turns out to be problematic. To see the nature of the problem, consider the representative neuroscientific approach to the problem of consciousness: to identify the neural correlates of consciousness (NCC). Content-specific NCC is usually defined as “the minimum neuronal mechanisms jointly sufficient for any one specific conscious percept” [60]. This definition is vague as to whether ‘neuronal mechanisms’ mean the anatomical structure and/or the activity states of the neurons in the relevant mechanisms. Typically, the anatomical structure is assumed to be of those that are usually found in the healthy brains of adult humans who can introspectively report their contents of consciousness with an accuracy. Under such an anatomical assumption, a pattern of neural activity in some specific anatomical location over a certain temporal period is usually considered as a content-specific NCC. An example of a face-related NCC is the extended neural activation in fusiform gyrus in the right hemisphere [1, 12, 57, 100]. If the activity in this area is transiently lost due to electrical stimulation, face perception gets disrupted without affecting other types of percept [85].

A traditional NCC approach can be described as a research paradigm, where a snapshot of pattern of neural activity N is minimally sufficient for a specific conscious phenomenology P. For example, Chalmers (2000) wrote, as one potential way to define the NCC for an arbitrary phenomenal property P, as follows:

“A neural correlate of a phenomenal family S is a neural system N such that the state of N directly correlates with the subject’s phenomenal property in S.”

From the CT perspective, this approach can be rephrased as the following. First, it tries to identify the neuronal category N as the category whose objects are patterns of neuronal activities in a specific region of the brain and whose arrows are transitional relationships between the patterns of neuronal activities. The phenomenal category P can be considered with its objects contents of consciousness with arrows transitional relationships. Second, it tries to find a sufficiently strong correlational relationship between the regions’ neuronal activities, e.g., its neuronal state, and the category of consciousness, regarding it as the NCC.

While this approach seems quite natural, it has several difficulties, as pointed out by others (e.g., see [18]). One of the fundamental issues is that neural activity pattern N needs to be defined within some anatomical reference frame. For example, face perception is typically correlated with the neural activity in fusiform face area (FFA) in normal healthy subjects. However, brain-damaged patients, whose damage spares more or less normal level of neural activity in FFA, can be impaired in face perception [86]. Considering even more extreme cases, almost nobody would argue that a conscious face phenomenology, p, arises from neural activity pattern N within FFA, which are artificially cut from the rest of the brain and kept alive and functional in a jar. Even if such an entity were to experience consciousness, unlike normal healthy humans, it would not experience it as a visual face phenomenology because it doesn’t have any capabilities to experience other possible phenomenologies to compare with [10].

To sum, most traditional NCC approaches implicitly require that the NCC to be embedded in some anatomical reference frame that extends beyond a single region as the NCC. That entails that the neuronal activities of two, if not more, regions will serve as the NCC which renders problematic the assumption of a single neuronal state N serving as the NCC. We must consequently raise the question for the exact relationship between the anatomical reference frame, e.g., different regions, and the neural activity patterns, e.g., the neuronal states. In this paper, to address this issue, we propose to consider the “relationship” between at least two neuronal categories, N0 and N1, instead of one single category N. In short, we consider N0 as the actual state of the neural activity of the actual network, and N1 as all possible states of the neural activity of all possible networks. As we argue, considering how the actual network state and structure is embedded in a larger possible network states and structures will allow us to clarify why we need to consider the anatomical reference frame to consider the NCC. This allows us to reconsider the relationship between the neural and the phenomenal category in a more nuanced way.

In the next section we will give more detailed explanation on the way to conceive how the two categories N0 and N1 are related to the phenomenal category P, in the context of IIT and TTC.

### Categories in IIT and TTC

Both IIT and TTC conceive a more complex notion of content-specific NCC that extends and goes beyond the assumption of a single neuronal state, e.g., category N, serving as content-specific NCC. They thus agree in that we need to introduce at least two neuronal categories (e.g., N0 and N1) to explain content-specific NCC. However, IIT and TTC differ in the exact formulation of the two neuronal categories. Note that we here do not go into details of both IIT and TTC; we focus on those aspects that are relevant within the present Category-Theoretical approach.

### Categories in IIT

For the full description of IIT, see [78, 101, 102]. Briefly, IIT starts from identifying the essential properties of phenomenologies (existence, composition, information, integration and exclusion, [78]) and then claim phenomenology is “identical” to the conceptual structures. Then IIT proposes several postulates based on which what types of physical mechanisms could potentially support such conceptual structures.

One essential aspect of IIT is that rather than focusing on only the actual state of a set of neurons, it considers the relationship between all possible states and an actual state of the set. Any conscious experience is informative in the sense that it specifies one of many *possible* experiences. Further, IIT considers how a system (or a mechanism) is potentially affected when the system is disconnected in all possible ways. In other words, IIT considers the relationship between all possible network configurations and an actual network configuration.

The original IIT can be regarded to propose a relationship between conscious experience (or phenomenal category P), conceptual structure (or informational structure category, I), and physical substrates (or neural category N), where P is “identical” to I [78, 102]. One way to view IIT is a functor (as we introduce in Functor, natural transformation and consciousness) from N to I (see Functors and natural transformations in IIT and TTC). So far, IIT just assumes that I is “identical” to P. Future work is needed to investigate the detailed formulation/analysis on the structure of I or P. Thus, in this paper, we will focus on how IIT treats the category of N through an *IIT functor* and a possible *IIT natural transformation* (as we introduce in Functor, natural transformation and consciousness) and demonstrate that a rigorous, yet complex, operations of IIT [78] can be re-interpreted through CT, which eventually offers a fresh and interesting insight which is not normally emphasised in IIT papers.

Let us start considering what are the essential categories in IIT and what corresponds to objects and arrows in category N0 and N1 in IIT^1^. We propose that in IIT an object is a stochastic causal network with transition probability matrix (TPM) to describe its state transition and an arrow is a manipulation on the network with the TPM (Fig 2). IIT considers various rules for the types of manipulations and selections of arrows. However, these manipulations can be relaxed or compared with various other types which will be an informative research direction in its own. For our purpose, it is important to note that IIT considers configurations of causal relationships by quantifying how each powerset of mechanism contributes to the whole. IIT does this by introducing an arrow that we call *decomposition.* Decomposition operation can be considered as something similar to marginalization. The purpose of the decomposition operation is to consider and quantify how much a neuron A contributes to a system of neurons A and B. Thus, given an object AB, we have at least three arrows: AB->AB (identity), AB->A and AB->B. Decomposition arrows capture one of the central properties of IIT: axiom and postulate of “composition” in consciousness.

**Fig 2.**
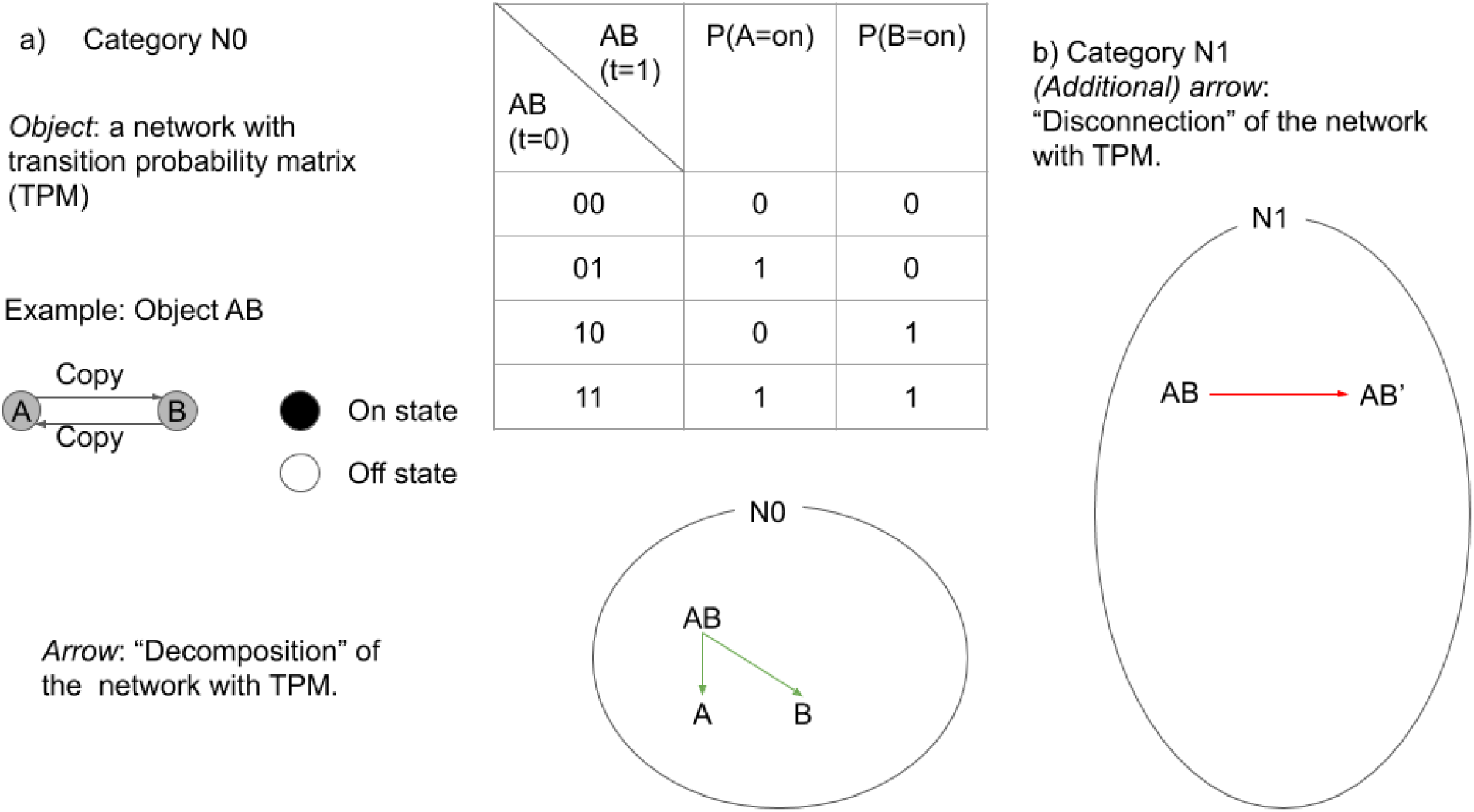
**a)** In IIT, category is defined by an object that is a stochastic network with transition probability matrix (TPM). The exemplar network is composed of a copy gate A and B, who copies the state of the other gate with a time delay of 1. The state of the gate is either on or off. The table on the right describes its TPM. An arrow in category N0 is “decomposition” of the network with TPM. Decomposition allows IIT to quantify the causal contribution of a part of the system to the whole. **b)** Disconnection arrows finds the minimally disconnected network, which capture concept of the amount of integration in IIT.

Note that N0 satisfies all the requirement to be a category (identity, associativity, and compositionality are all satisfied).

Next, consider a category, N1, in which objects are *all possible* networks associated with TPM. N1’s arrows are *decomposition* as in N0 and also *disconnection*. Disconnection operation can be considered as transformation of TPM to another TPM through (virtual) disconnection of the network, such that subsets of the network are statistically independent [79, 99]. Disconnections arrows capture another central property of IIT: axiom and postulate of “integration” in consciousness. The disconnection arrow can be related to the amount of integrated information.

Again, note that N1 also satisfies all the requirement to be a category. In Functor, natural transformation and consciousness, we will discuss how these categories are related through functors.

### Categories in TTC

Unlike IIT, the TTC does not consider different neurons’ or regions’ activities as starting point to distinguish different neuronal states. Instead, TTC stresses the temporal dimension and thus the dynamics of neuronal activity as it operates across different regions and points in time (for details, see [REF X, Y, Z]). Specifically, for the TTC, N0 and N1 are both the temporal dynamics of neural systems (extending possibly across all brain areas). N0 precedes N1 in time. In other words, N0 can be regarded as pre-stimulus (which ultimately can be traced to the continuously ongoing dynamics of the spontaneous activity) and N1 as post-stimulus neural activity.

To support this claim, we consider the empirical data in both fMRI [14, 45, 88, 89] and EEG/MEG [2, 8, 11, 66], which show that the amplitude and/or variance of pre-stimulus activity plays a major role in whether the subsequent stimulus and its respective contents becomes conscious or not. Typically, high pre-stimulus activity levels, e.g., high amplitude or variance, are more likely to allow for associating contents with consciousness than low pre-stimulus activity levels. Baria et al. (2017) showed that the pre-stimulus activity level at up to 1.8s prior to stimulus onset can predict (on a single trial level; above chance) whether a visual content will be consciously seen or not. Moreover, pre-stimulus activity levels are not only relevant in the region typically processing the respective stimulus, e.g., like FFA for face stimuli, auditory cortex for auditory stimuli, etc. Additionally, the pre-stimulus activity level in other more distant regions like parietal and prefrontal cortex have also been shown to be relevant in impacting conscious perception of an object during the post-stimulus period [14, 45, 88, 89].

Together, these data suggest that both pre-stimulus activity levels, e.g., amplitude and/or variance, and post-stimulus activity level may need to be included in content-specific NCC. Specifically, as emphasized by TTC, it is the temporal and spatial dynamics of the pre-stimulus activity, e.g., its variance being present in different regions, that is central for associating post-stimulus activity and its contents with consciousness (see below for more details on the pre-post-stimulus dynamics and how it allows for a particular visual stimulus to be consciously perceived).

Accordingly, the TTC entails a more complex notion of content-specific NCC that extends and goes beyond a single neuronal state (and thus also beyond the neural prerequisites of consciousness; [4, 22]) when assuming the temporo-spatial dynamics of two distinct neuronal states to underlie consciousness. Mathematically, that requires two distinct neuronal categories, i.e., N0 and N1 to formalize content-specific NCC of TTC within the context of CT.

To be more explicit, for TTC, objects of N0 and N1 are neural activity over time and space, and arrows are explicitly defined only for identity. This guarantees that N0 and N1 are both category.

In sum, as a critical component to consider consciousness, IIT considers all possible states (N1) and an actual state (N0) of a mechanism, while TTC considers temporal dynamics of pre-(N0) and post-(N1) stimulus neural activities as objects of these categories. Although the specifics are different, both are clearly going beyond a traditional NCC conceptualization; a particular neural state at a given time N to correspond to a particular phenomenal state P. Rather, they both point that it is a “relationship” between N0 and N1 that corresponds to a particular phenomenal state P. In the next section, we will introduce a mathematical tool to consider a relationship between the two categories; functor.

## Functor, natural transformation and consciousness

### Definition of functor and natural transformation

A functor is defined as a structure-preserving transformation between two categories. In fact, a functor is defined as an arrow in “the category of categories”, shown in Fig 3 below.

**Fig 3.**
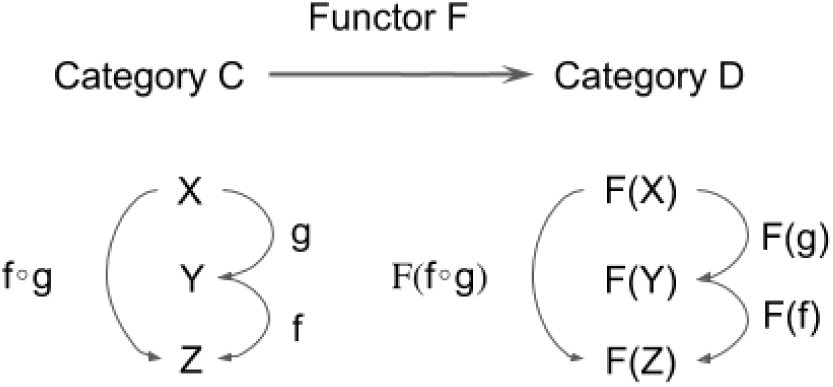
Definition 3. A correspondence F from a category C to another category D which maps each object/arrow in C to corresponding object/arrow in D is called a functor if it satisfies the following 3 conditions: 1. It maps f : X → Y in C to F(f) : F(X) → F(Y) in D. 2. F(f∘g) = F(f)∘F(g) for any (composable) pair of f and g in C. 3. For each X in C, F(1X)=1F(X).

In short, a functor is a correspondence which preserves categorical structure. Through a functor, one category and its associated structure is related to those in another category and its associated structure. A functor allows us to consider a possibility to relate obviously different domains (e.g., the phenomenal and the neuronal) to each other.

One of the most important notions that we introduce in the present paper is what we describe as an “inclusion functor” (Fig 4a). Let us consider two categories *C* and *D.* A functor F from C to D is called an inclusion functor if

**Fig 4.**
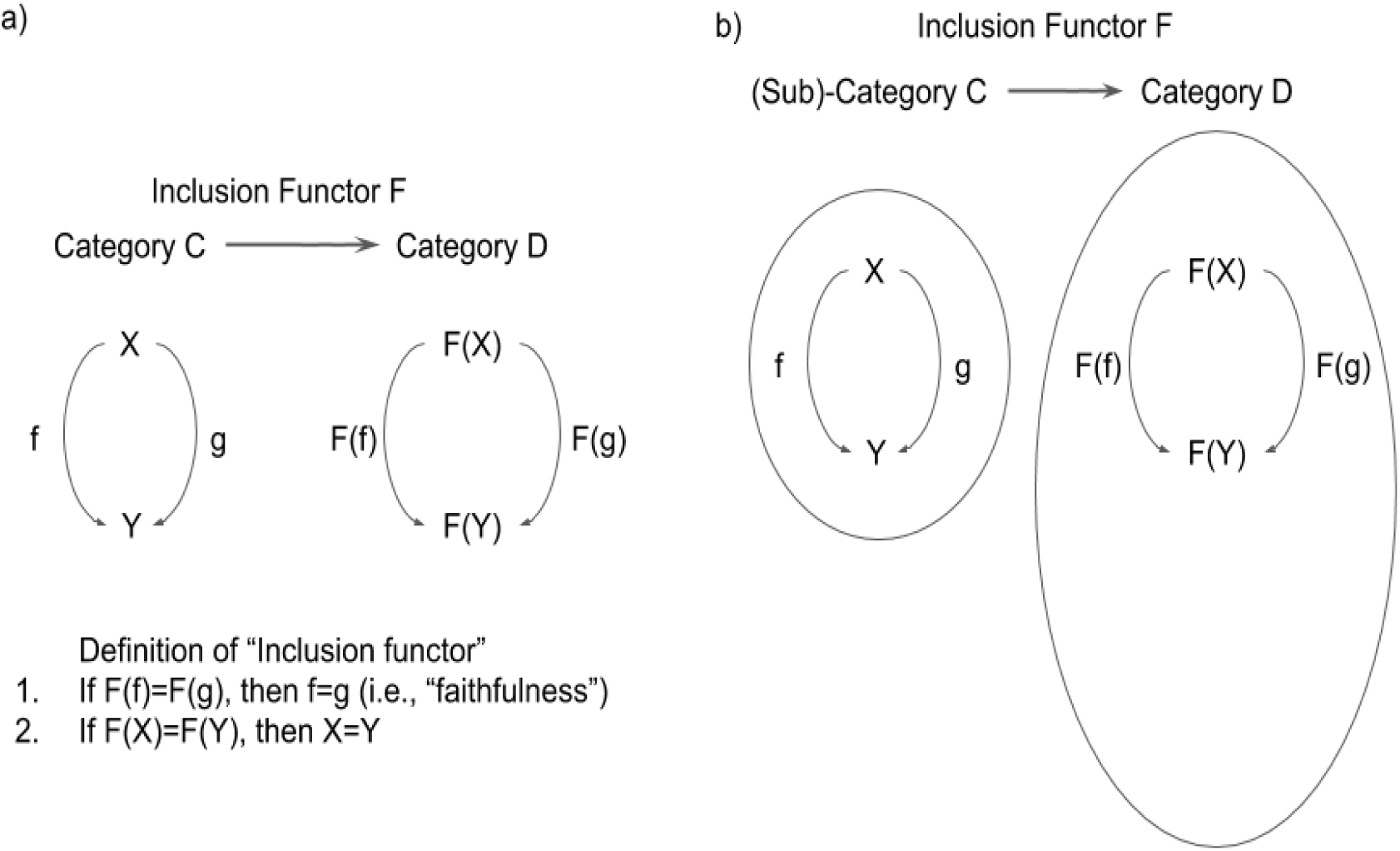
**a)** Definition of “inclusion functor”. **b)** Sub-category C is included by Category D if Inclusion Functor F: C->D exists. Note that C does not not need to be “a part of” D to be “included” (unlike a commonsense definition of “inclusion”).

1. For any pair of object X, Y in *C* and arrows f, g in *C* from X to Y, i.e. dom(f)=dom(g)=X and cod(f)=cod(g)=Y, F(f)=F(g) implies f=g. (Functors satisfying this condition is called “faithful’’.)
2. For any pair of objects X and Y in *C*, F(X)=F(Y) implies X=Y.

When there is an inclusion functor from *C* to *D, C* is called as a subcategory of *D*.^2^

The intuition for the terms can be explained as follows (also see Fig 2b): Let us consider the situation that any object/arrow in *C* has a corresponding object/arrow in *D*, and the notion of dom/cod, composition and identity for *C* are in common with those for *D*. Then it is quite natural to think of *C* as subsystem of *D* and thus to call *C* a subcategory of *D.* In this situation, we can define an inclusion functor F as a map sending each object/arrow in *C* as object/arrow in *D*, i.e. F(X)=X and F(f)=f for any object X and arrow f (Here X and f in the left hand side are an object and an arrow in *C* and those in the right hand side are those considered in *D*).

Let us briefly summarize the meaning of the inclusion functor. The existence of the inclusion functor from N0 to N1 essentially means that N0 is a “subcategory” of N1. Here we propose an inclusion functor i, which plays a fundamental role in this paper. i is defined by i(X)=X and i(f)=f (Fig 5). It works as the “basis” of the consciousness phenomena.

**Fig 5.**
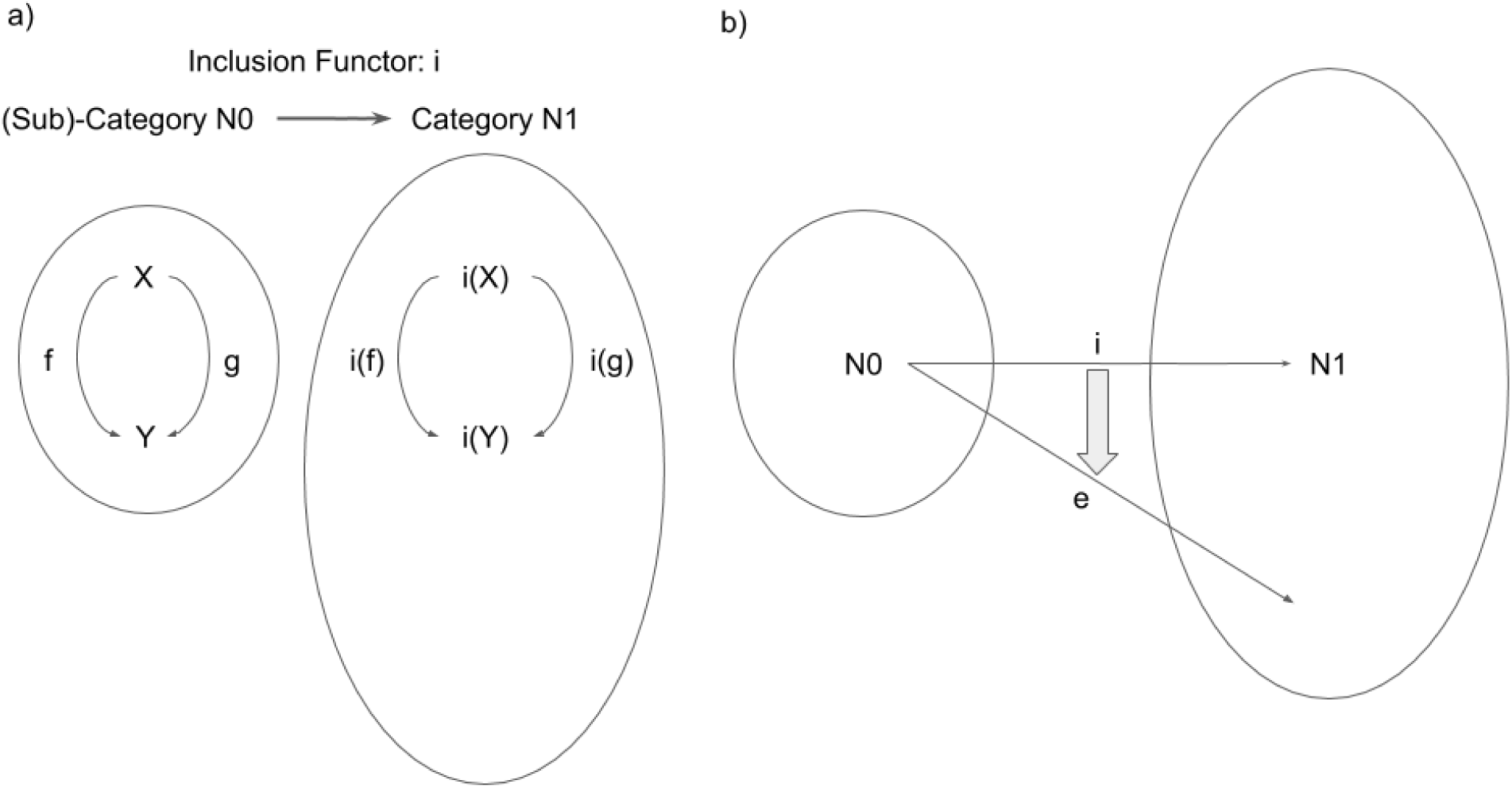
**a)** Inclusion Functor i: N0->N1. N0 is included in N1 through Inclusion Functor i. **b)** Expansion Functor e:N0->N1. e is a different structure preserving mapping from N0 to N1 (i.e., a functor from N0 to N1), but there is “natural transformation” from i to e.

To define the notion of expansion functor, which is a functor different from inclusion functor but closely related to it, we need to define the notion of “natural transformation” as relation between functors.

A functor is an “arrow” between two categories, but a functor can be can also be considered as an “object” in CT (as an arrow can be considered as an object. See Definition of category). When we consider functors themselves as “objects”, we call “arrows” between functors as “natural transformations”.

The definition of natural transformations is the following (Fig 6):

**Fig 6.**
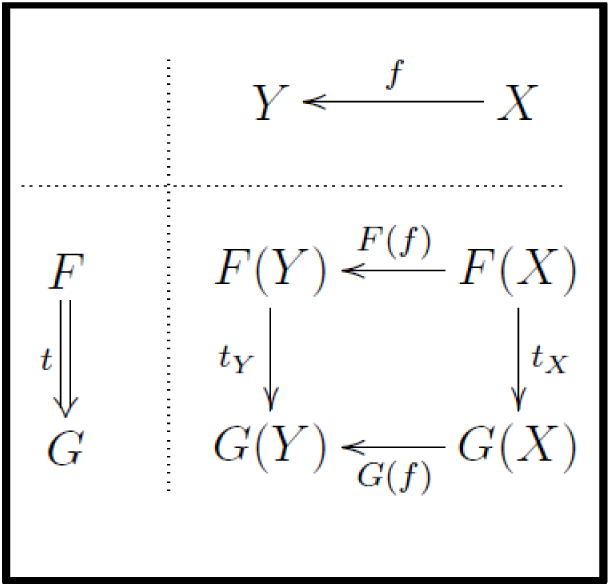
Definition 4. Let F, G be functors from category C to category D, a correspondence t is called a natural transformation from F to G if it satisfies the following two conditions: 1. t maps each object X in C to corresponding arrow tX: F(X) → G(X) in D. 2. For any f : X → Y in C, tY∘F(f) = G(f)∘tX.

For the natural transformation we use the notation such as t: F ⇒ G. The second condition above is depicted in Fig 6:

Upper-right part denotes the arrow in C *(f : X* → *Y)* and lower-left part the natural transformation from F to G (t: F ⇒ G). The second condition in the definition of natural transformation means that the diagram in the lower-right part commutes.

Intuitively speaking, *a natural transformation from functor F to functor G is the system of arrows indexed by objects*, which satisfies certain consistency condition. This is an interesting property of CT, which is worth emphasizing. A meta-level and abstract concept of a natural transformation is represented as a set of lower-level and concrete concept of arrows in a category. (We believe this nested mathematical structure of CT is particularly suited to capture some aspects of consciousness, which we will describe elsewhere.)

Now, equipped with this notion of a natural transformation, we can talk about a “structure preserving map” between two functors. Now we introduce the notion of an “expansion functor”, as a functor toward which there is a natural transformation from inclusion functor^3^. That is, an expansion functor, e, is an expanded form, or a version of the inclusion functor, i, transformed through some natural transformation.

### Functor, natural transformation and consciousness

With the concepts of inclusion and expansion functors and natural transformation, we can now propose to provide a more explicit relationship between the neural activity N to the anatomical structure where N is embedded, which is a necessary step to go beyond the traditional NCC approach. In the present paper we use the inclusion functor i as the basis and expansion functors as its expanded version, to stress the importance of the idea that expansion functors are the variations from the functor i as the basis through some natural transformation.

In the next section, we point out that some essential aspects of both IIT and TTC can be captured by a consideration of different versions of expanding functors generated from the inclusion functor. We will see that both IIT and TTC distinguish between inclusion and expansion functor. As in the case of N0 and N1, interestingly, we will see that both IIT and TTC implicitly incorporate inclusion functors, expansion functors and natural transformations between them in different manners. Regardless of the specifics of the theories, we argue that these concepts of functor and natural transformation are one of the missing components of the traditional NCC research.

## Functors and natural transformations in IIT and TTC

### Functors and natural transformations in IIT

Let us first reinterpret some aspects of IIT in CT, especially with the concepts of inclusion functor, expansion functors and natural transformations between them. A critical concept in IIT, the amount of integrated information, *φ* (small phi), can be interpreted as the “difference” between the actual and the (minimally) disconnected network [79, 80, 99]. This can be captured by CT concepts of inclusion and expansion functors. The compositional aspects of IIT, or a set of *φ*s corresponds to a set of objects captured by these functors. The big phi, *Φ*, which corresponds to quantity (e.g., level) of consciousness, or system level integration, can now be interpreted as a natural transformation.

Let us unpack the above statements. As we explained in Fig 1 in Categories in IIT and TTC, we consider a category N0, to be composed of objects (the actual network with TPM) and arrows, which “decompose” the system. We also consider another category N1, composed of all possible networks with TPM and arrows. In addition to “decomposition” arrows, N1 is equipped with “disconnection” arrows. Obviously, N0 is “included” by N1. Inclusion functor, *i*, finds the corresponding objects and arrows in N1.

Now, we define an expansion functor, *e*, as the one that finds the minimally disconnected version of the original network in N1 (Fig 7a).

**Fig 7.**
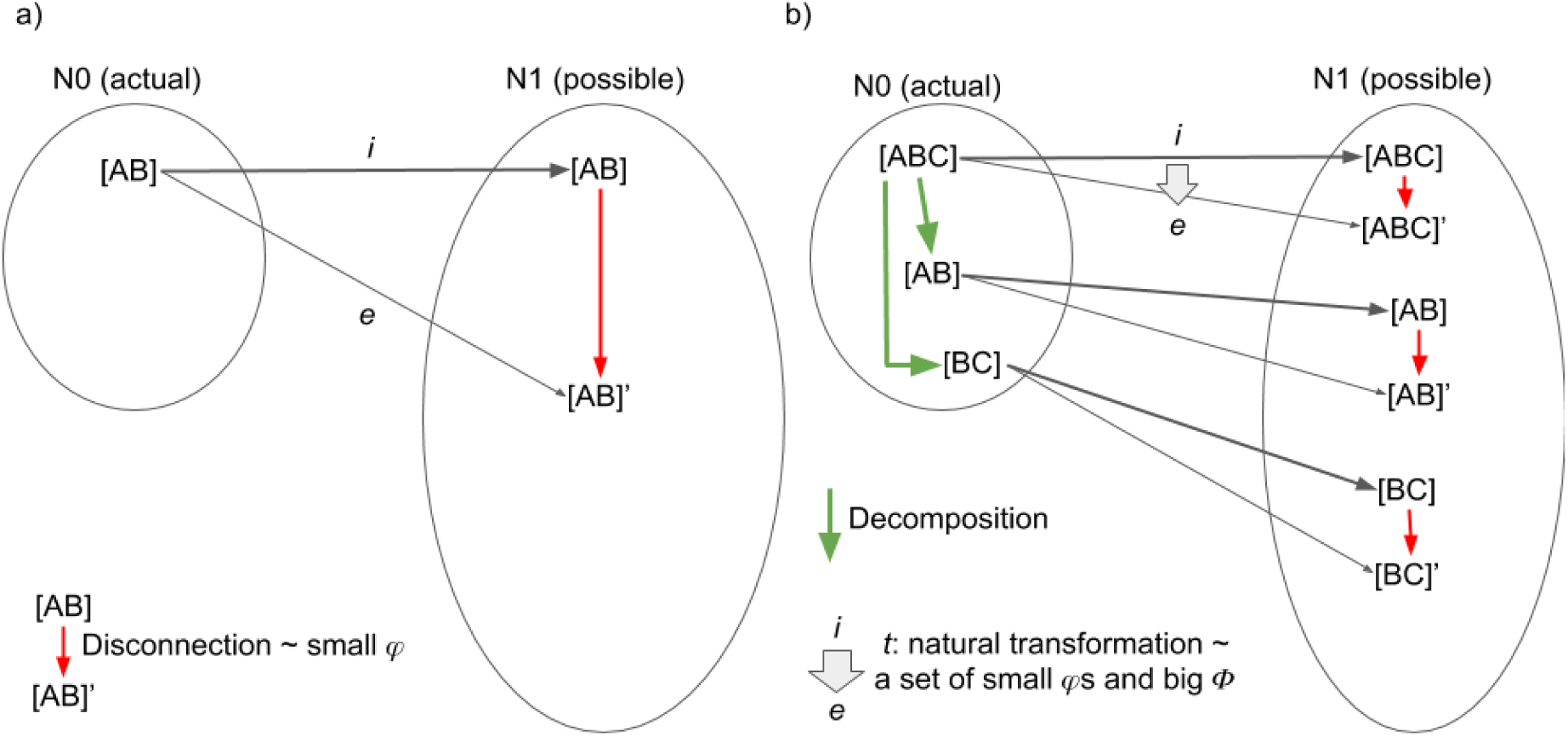
**a)** Inclusion functor, i, expansion functor, e, in the IIT category N0 (actual) and N1 (all possible). Objects in N0 and N1 (e.g., [AB]) are a network with TPM, and arrows in N0 and N1 are manipulation of network/TPM that is allowed in IIT. Within N0, we consider only decomposition arrows. N1 is enriched by additional disconnection arrows that represent an operation that finds a “minimally disconnected” network with TPM within N1. An expansion functor, e, finds the minimally disconnected network (e.g., [AB]’) of the original network (e.g., [AB]). e also preserves the structure of N0, and qualifies as a functor. A red arrow within N1 that goes from the actual to the minimally disconnected network corresponds to integrated information, φ. **b)** Considering decomposition arrows in N0 allows N0 to consist of a powerset of the network. If natural transformation, t, from the inclusion to the expansion functor exists, t gives us a power set of φ’s, the original and the minimally disconnected network with TPMs. This corresponds to system level integration, Φ.

Together with the inclusion functor, the expansion functor from the original network and TPM now gives us a set of *φ*’s. Not only the original network (e.g., ABC) but also its subnetwork components (e.g., AB and BC) have corresponding *φ*, which is derived by corresponding disconnection arrows in N1.

Now, we assume there is a natural transformation between inclusion and expansion functors. Then, a set of *φ*’s are obtained by a natural transformation, *t*, between the inclusion and the expansion functors. This set can quantify integration at the system level, which corresponds to what IIT calls *Φ* (big phi). The concept of natural transformation clarifies the essence of IIT; IIT is a theory that proposes a set of *φ*’s and *Φ*, which corresponds to quality (e.g., qualia, contents) and quantity (e.g., level) of consciousness, respectively.

Does a natural transformation, t, really exist? Let’s consider it in Fig 8. If t qualifies as a natural transformation, e(f), that is, a decomposition arrow from AB to A in N0 (or N1) has to correspond to a decomposition arrow in N1 from AB’ and A’. As far as we know (including our personal communication with Dr. Masafumi Oizumi), IIT has never considered a precise mathematical formulation between the disconnected networks. Thus, while we know that some kind of relationship exists between e(AB) and e(A), at this point, we do not know if the operations that are used to “decompose” AB into A can (i.e., f) be directly applied to the “disconnected” AB’ into A’.

**Fig 8.**
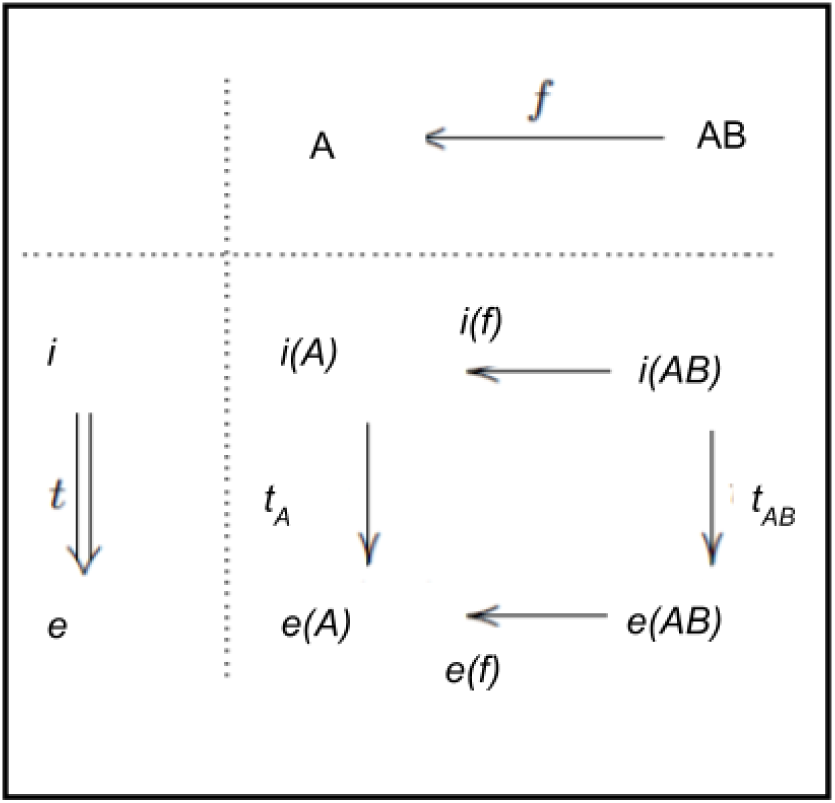
Natural transformation, t.

Here, let us briefly remark a potential consequence of the existence of a natural transformation. If one can describe the “decomposition” arrow between the disconnected networks in a formal mathematical relationship, which parallels the “decomposition” arrow between the original networks, then we can prove the existence of a natural transformation between inclusion and expansion functor. Mathematically, this guarantees a possibility of building up a larger network by considering a larger context (say, adding C into AB) in IIT. IIT papers, according to our understanding, has been so far mute on a possibility or limitation of this “reverse-reductionism” approach. Intuitively, however, the role of AB among ABC should be similar to the role of AB among ABCD (to some extent). Our preliminary results indeed suggest this may be the case, when integrated information is computed from the neural recording data (Leung et al 2018 ACNS poster, in preparation). If it is indeed the case that we can reverse-reductionistically understand the whole by building up and pasting many parts of the systems (potentially using presheaf theory [REF]), then this approach may make IIT more mathematically tractable.

On the other hand, it is totally possible that there is no formal arrow like e(f). If that is the case, it practically means that the integrated information of a part of the system can completely and unpredictably change based on the way it is embedded in the context. This may reveal an extreme holistic property of the IIT. Given the phenomenological axiom of compositionality in IIT, however, we surmise that such a result probably requires a revision of the postulate of the IIT.

In summary, this conjecture, a necessity and potential consequence of consideration between the disconnected networks, is a direct consequence of considering IIT from the CT perspective, which may prove useful in the future mathematical examination of IIT.

In sum, the category theoretic reinterpretation of IIT tells us that to construct quantitative theory of consciousness, consideration of the relation between actual and possible is necessary: expanding functor e (as a mapping towards a set of disconnected networks), or more precisely,” expanding functor in relation to inclusion functor”. In terms of category theory, natural transformation from i to e provides us a set of small phi, integrated information, which characterize quality of consciousness, and a big phi, the system level integration or quality of consciousness. Quality and quantity of consciousness in IIT amounts to the quantitative evaluation on the natural transformation from i to e (if it exists). As we have defined, a natural transformation is nothing but a system of arrows indexed by objects which satisfies certain consistency condition, which requires further investigation.

### Functor and natural transformations in TTC

Now, TTC is well compatible with the concepts of inclusion and expansion functors and a natural transformation as TTC also emphasizes the need for conceiving the relationship between pre-stimulus activity (N0) and post-stimulus activity^4^ (N1) in terms of integration (but not in the sense used in IIT). However, unlike IIT, TTC again emphasizes the dynamic, e.g., temporo-spatial mechanisms that are supposedly underlying the relationship between pre- and post-stimulus activity including their integration.

To be more specific, an inclusion functor, i, from N0 to N1 would correspond to a mapping from the pre-stimulus to post-stimulus neural activity without sensory input (or any other perturbation). An expansion functor, e, would correspond to a mapping from the pre-stimulus to post-stimulus neural activity with a specific sensory input (or any other perturbation). Expansion functors, therefore, are a family of functors. Natural transformation between i and e describes relationships among all possible consequences of perturbations.

Traditional models presuppose that stimulus-induced activity as related to external stimuli is simply added to and thus supervenes on the ongoing internal neuronal activity – this amounts to additive rest-stimulus interaction [3, 6, 34, 35, 36, 96] In contrast, recent findings suggest non-additive interaction between pre- and post-stimulus activity levels as based on EEG [43], fMRI [31, 32, 50, 88],and computational modelling [84].

In the case of non-additive interaction, the post-stimulus activity is not simply added on or supervenes upon the pre-stimulus activity level. Instead, the level of pre-stimulus activity exerts a strong impact on the level of subsequent post-stimulus activity. In terms of the response amplitude, low pre-stimulus activity levels lead to relative higher post-stimulus activity levels than high pre-stimulus activity levels [32, 50, 51]. Importantly, recent studies in both MEG [11, 66] and fMRI [51] demonstrate that pre-stimulus variance and its non-additive impact on post-stimulus amplitude/variance is related to conscious contents [2, 11, 66, 93] and the level/state of consciousness [51]. Most interestingly, a recent study demonstrated that pre-post-stimulus variance are accompanied by the Lempel-Zev Complexity (LZC) in the pre-stimulus interval [107, 108]. IIT inspired a measure of level of consciousness, called Perturbational Complexity Index, combining a TMS-EEG experiment with the LZC [13]. While how integrated information relates PCI is unclear at this point, it raises a possible link between the non-additive dynamics of pre-post-stimulus interaction, as pointed outed in TTC, with information integration in IIT.

Another point on the importance of inclusion functor, exclusion functor and natural transformation between them in the context of TTC is the importance of N1 (post-stimulus activity) in relation to N0 (pre-stimulus activity). As the reviewed empirical evidence suggests, post-stimulus activity (N1), reflecting the processing of the contents themselves, is not sufficient to explain any particular phenomenology, p, on its own. In addition to post-stimulus activity (N1), pre-stimulus activity (N0) and its dynamics is necessary as N0 strongly affects and modulates how the subsequent N1 is processed. As such, TTC claims “consciousness does not come with the contents themselves” [75, chapter 6]. Instead, TTC suggests that consciousness is associated to the contents rather than coming with the contents themselves [72, 77]. Empirically, this means that the focus shifts from the neural activity in post-stimulus period itself to pre-stimulus activity and how it interacts with the stimulus, e.g., the non-additive dynamics of pre-post-stimulus interaction. Mathematically, that very same dynamics of non-additive pre-post-stimulus interaction can be well formalized by the here assumed natural transformation from inclusion functor to expansion functor.

## Conclusion

We here introduced Category Theory (CT) to account and formalize the relationship between the neuronal (N0 and N1) and phenomenal (P) domains in the neuroscience of consciousness. Specifically, we introduced four fundamental concepts in CT (category, inclusion, expansion functors, and natural transformation) in the context of the two major neuroscientific theories of consciousness, e.g., IIT and TTC. Now we briefly review some major implications for our search of the NCC in general in the future neuroscientific studies of consciousness.

The first point we made in this paper was that we need to distinguish between two different neuronal categories, N0 and N1, which both IIT and TTC implicitly have proposed. This approach seems to solve a difficulty in the traditional NCC research, which implicitly assumes the anatomical frame when it considers one specific neuronal state and its corresponding one specific phenomenal state. By explicitly considering two neural categories, both IIT and TTS consider N0 (a particular neural state) as embedded with N1 (all possible states), which is constrained by the anatomical reference frame.

The second point we made was even more important; a shift of the focus from the relationship between neuronal and phenomenal states, as promoted by the traditional NCC approach, to the relationship and, specifically, a particular form of relationship or interaction between two neuronal category (N0 and N1) as central for yielding consciousness. This emphasis of the relationship can be framed as natural transformation between inclusion and expansion functors. Addressing the same question, IIT and TTC provide different answers, e.g., a set of integrated information (IIT) and non-additive interaction (TTC).

Taken together, we conclude that CT provides the mathematical tools to formalize the relationship between the neuronal and the phenomenal domains and to give a blueprint on how to extend it beyond the traditional NCC approaches. In the context of IIT, mathematical investigation on the existence of natural transformation between inclusion and expansion functors can be potentially fruitful investigation, as it may allow “reverse reductionistic” approach to understand a large network to overcome the fundamental difficulty in IIT. In the context of TTC, CT can extend the concept of non-additivity into temporo-spatial dynamics.

As such, we conclude that the introduction of CT in the study of neural correlates of consciousness awaits further fruitful theoretical development, with its potential to connect or translate across different theories of consciousness, which we could not mention in this paper (e.g., the Global Neuronal Workspace Theory (GNWT) [23-25] higher order theories of consciousness [63, 87], recurrent processing theory [62], operational space and time [33], neural synchrony [30], and social and attention schema theory [37]). Comparison of the theories through CT, as we did for IIT and TTC here, may inspire development of an entirely novel approach to connect neuronal and phenomenal domains in a formal and mathematical way.

## Acknowledgements

NT was funded by Australian Research Council Discovery Project grants (DP180104128, DP180100396), and a grant (TWCF0199) from Templeton World Charity Foundation, Inc. HS was partially supported JST CREST(JPMJCR17N2) and JSP-KAKENHI (JP17H01277). HS and NT acknowledge informal comments by Masafumi Oizumi. NT acknowledge informal discussion among Monash Neuroscience of Consciousness laboratory.

## Footnotes

1 Throughout, we only consider the state-independent measures of IIT [9] for simplicity, but generalization to the state-dependent measures of IIT [78] is not difficult.)

2 From a more radically category-theoretic viewpoint, the inclusion functor F itself is called subcategory, but in the present paper we call *C* as subcategory to avoid confusion.

3 Note that an “expanding functor” is not the standard term in CT: We name it for the importance in consciousness studies. On the other hand, “inclusion functor” is a standard term in mathematics.

4 or more precisely, all possible activities which include the post-stimulus activities.

## References

1. Allison T, Ginter H, McCarthy G, Nobre AC, Puce A, Luby M, Spencer DD. Face recognition in human extrastriate cortex. J Neurophysiol. 1994;71(2):821–5. doi: 10.1152/jn.1994.71.2.821

2. Arazi A, Censor N, Dinstein I. Neural Variability Quenching Predicts Individual Perceptual Abilities. J Neurosci. 2017;37(1):97–109. doi: 10.1523/JNEUROSCI.1671-16.2016.

3. Arieli A, Sterkin A, Grinvald A, Aertsen A. Dynamics of ongoing activity: explanation of the large variability in evoked cortical responses. Science. 1996;273(5283):1868–71.

4. Arzi-Gonczarowski Z. Perceive this as that – Analogies, artificial perception, and category theory. Ann Math Artif Intell. 1999;26(1-4):215–252. doi: 10.1023/A:101896302

5. Aru J, Bachmann T, Singer W, Melloni L. Distilling the neural correlates of consciousness. Neurosci Biobehav Rev. 2012;36(2):737–46. doi: 10.1016/j.neubiorev.2011.12.003.

6. Azouz R, Gray CM. Cellular mechanisms contributing to response variability of cortical neurons in vivo. J Neurosci. 1999;19(6):2209–23.

7. Baez JC, Stay M. Physics, Topology, Logic and Computation: A Rosetta Stone. https://arxiv.org/abs/0903.0340

8. Bai Y, Nakao T, Xu J, Qin P, Chaves P, Heinzel A, Duncan N, Lane T, Yen NS, Tsai SY, Northoff G. Resting state glutamate predicts elevated pre-stimulus alpha during self-relatedness: A combined EEG-MRS study on “rest-self overlap”. Soc Neurosci. 2011;11(3):249–63. doi: 10.1080/17470919.2015.1072582.

9. Balduzzi D, Tononi G. Integrated information in discrete dynamical systems: motivation and theoretical framework. PLoS Comput Biol. 2008;4(6):e1000091. doi: 10.1371/journal.pcbi.1000091.

10. Balduzzi D, Tononi G. Qualia: the geometry of integrated information. PLoS Comput Biol. 2009;5(8):e1000462. doi: 10.1371/journal.pcbi.1000462.

11. Baria AT, Maniscalco B, He BJ. Initial-state-dependent, robust, transient neural dynamics encode conscious visual perception. PLoS Comput Biol. 2017;13(11):e1005806. doi: 10.1371/journal.pcbi.1005806.

12. Baroni F, van Kempen J, Kawasaki H, Kovach CK, Oya H, Howard MA, et al. Intracranial markers of conscious face perception in humans. Neuroimage. 2017;162:322–343. doi: 10.1016/j.neuroimage.2017.08.074.

13. Bayne T. The unity of consciousness. Oxford: Oxford University Press; 2010.

14. Boly M, Phillips C, Tshibanda L, Vanhaudenhuyse A, Schabus M, Dang-Vu TT, et al. Intrinsic brain activity in altered states of consciousness: how conscious is the default mode of brain function? Ann N Y Acad Sci. 2008;1129:119–29. doi: 10.1196/annals.1417.015.

15. Casali AG, Gosseries O, Rosanova M, Boly M, Sarasso S, Casali KR, et al. A theoretically based index of consciousness independent of sensory processing and behavior. Sci Transl Med. 2013;5(198):198ra105. doi: 10.1126/scitranslmed.3006294.

16. Chalmers DJ. The conscious mind. Oxford:Oxford University Press; 1996.

17. Chalmers DJ. What is a neural correlate of consciousness? Neural correlates of consciousness: Empirical and conceptual questions. Cambridge: MIT Press; 2000.

18. Chialvo DR. Emergent complex neural dynamics. Nat Phys. 2010;6:744–750. doi:10.1038/nphys1803

19. Churchland P. Brain-wise. Cambridge:MIT Press; 2002.

20. Crick F, Koch C. Some reflections on visual awareness. Cold Spring Harb Symp Quant Biol. 1990;55:953–62.

21. Crick F, Koch C. A framework for consciousness. Nat Neurosci. 2003;6:119–126. doi:10.1038/nn0203-119

22. de Graaf TA, Hsieh PJ, Sack AT. The ‘correlates’ in neural correlates of consciousness. Neurosci Biobehav Rev. 2012;36(1):191–7. doi: 10.1016/j.neubiorev.2011.05.012.

23. Dehaene S, Charles L, King JR, Marti S. Toward a computational theory of conscious processing. Curr Opin Neurobiol. 2014;25:76–84. doi:10.1016/j.conb.2013.12.005

24. Dehaene S, Changeux JP. Experimental and theoretical approaches to conscious processing. Neuron. 2011;70:200–227. doi:10.1016/j.neuron.2011.03.018

25. Dehaene S, Naccache L. Towards a cognitive neuroscience of consciousness: basic evidence and a workspace framework. Cognition. 2001;79(1-2):1–37.

26. Ehresmann AC, Vanbremeersch JP. Hierarchical evolutive systems: a mathematical model for complex systems. Bull Math Biol. 1987;49(1):13–50.

27. Ehresmann AC, Vanbremeersch JP. Information processing and symmetry-breaking in memory evolutive systems. Biosystems. 1997;43(1):25–40.

28. Ehresmann AC, Gomez-Ramirez J. Conciliating neuroscience and phenomenology via category theory. Prog Biophys Mol Biol. 2015;119(3):347–59. doi: 10.1016/j.pbiomolbio.2015.07.004.

29. Eilenberg S, MacLane S. Relations between homology and homotopy groups of spaces. Ann Math. 1945;46:480–509. doi:10.2307/1969165.

30. Engel AK, Singer W. Temporal binding and the neural correlates of sensory awareness. Trends Cogn Sci. 2001;5(1):16–25.

31. Ferri F, Costantini M, Huang Z, Perrucci MG, Ferretti A, Romani GL, et al. Intertrial variability in the premotor cortex accounts for individual differences in peripersonal space. J Neurosci. 2015;35(50):16328–39. doi: 10.1523/JNEUROSCI.1696-15.2015.

32. Ferri F, Nikolova YS, Perrucci MG, Costantini M, Ferretti A, Gatta V, et al. A Neural “Tuning Curve” for Multisensory Experience and Cognitive-Perceptual Schizotypy. Schizophr Bull. 2017;43(4):801–813. doi: 10.1093/schbul/sbw174.

33. Fingelkurts, AA, Fingelkurts, AA, Neves CF. Natural world physical, brain operational, and mind phenomenal space-time. Phys Life Rev, 2010;7(2):195–249. doi:10.1016/j.plrev.2010.04.001

34. Fox MD, Snyder AZ, Zacks JM, Raichle ME. Coherent spontaneous activity accounts for trial-to-trial variability in human evoked brain responses. Nat Neurosci. 2006;9(1):23–5.

35. Fox MD, Snyder AZ, Vincent JL, Raichle ME. Intrinsic fluctuations within cortical systems account for intertrial variability in human behavior. Neuron. 2007;56(1):171–84. doi: 10.1016/j.neuron.2007.08.023

36. Fox MD, Raichle ME. Spontaneous fluctuations in brain activity observed with functional magnetic resonance imaging. Nat Rev Neurosci. 2007;8(9):700–11.

37. Graziano MS, Kastner S. Human consciousness and its relationship to social neuroscience: A novel hypothesis. Cogn Neurosci. 2011;2(2):98–113. doi: 10.1080/17588928.2011.565121

38. Hardstone R, Poil SS, Schiavone G, Jansen R, Nikulin VV, Mansvelder HD, et al. Detrended fluctuation analysis: a scale-free view on neuronal oscillations. Front Physiol. 2012;3:450. doi: 10.3389/fphys.2012.00450.

39. He BJ, Zempel JM, Snyder AZ, Raichle ME. The temporal structures and functional significance of scale-free brain activity. Neuron. 2010;66:353–369. doi: 10.1016/j.neuron.2010.04.020.

40. He BJ. Scale-free properties of the functional magnetic resonance imaging signal during rest and task. J Neurosci. 2011;31(39):13786–13795. doi: 10.1523/JNEUROSCI.2111-11.2011

41. He BJ. Scale-free brain activity: past, present, and future. Trends Cogn Sci. 2014;18(9);480–7. doi:10.1016/j.tics.2014.04.003

42. He BJ. Spontaneous and task-evoked brain activity negatively interact. J Neurosci. 2013;33:4672–4682. doi:10.1523/JNEUROSCI.2922-12.2013

43. He BJ, Zempel JM. Average is optimal: an inverted-U relationship between trial-to-trial brain activity and behavioral performance. PLoS Comput Biol. 2013;9(11):e1003348. doi: 10.1371/journal.pcbi.1003348.

44. Healy MJ, Caudell TP, Goldsmith TE. A Model of Human Categorization and Similarity Based Upon Category Theory. Albuquerque (NM): University of New Mexico, Electrical & Computer Engineering Technical Reports; 2008 Jul. Report No.: EECE-TR-08-0010,.

45. Hesselmann G, Kell CA, Eger E, Kleinschmidt A. Spontaneous local variations in ongoing neural activity bias perceptual decisions. Proc Natl Acad Sci USA. 2008;105(31):10984–9. doi: 10.1073/pnas.0712043105.

46. Hoel EP, Albantakis L, Tononi G. Quantifying causal emergence shows that macro can beat micro. Proc Natl Acad Sci USA. 2013;110(49):19790–5. doi: 10.1073/pnas.1314922110.

47. Honey CJ, Thesen T, Donner TH, Silbert LJ, Carlson CE, Devinsky O, et al. Slow cortical dynamics and the accumulation of information over long timescales. Neuron. 2012;76(2):423–34. doi: 10.1016/j.neuron.2012.08.011.

48. Poincare H. Science and Method. London: Nelson Publisher; 1908.

49. Huang Z, Zhang J, Wu J, Qin P, Wu X, Wang Z, et al. Decoupled temporal variability and signal synchronization of spontaneous brain activity in loss of consciousness: An fMRI study in anesthesia. Neuroimage. 2016;124(Pt A):693–703. doi: 10.1016/j.neuroimage.2015.08.062.

50. Huang Z, Zhang J, Longtin A, Dumont G, Duncan NW, Pokorny J, et al. Is There a Nonadditive Interaction Between Spontaneous and Evoked Activity? Phase-Dependence and Its Relation to the Temporal Structure of Scale-Free Brain Activity. Cereb Cortex. 2017;27(2):1037–1059. doi: 10.1093/cercor/bhv288.

51. Huang Z, Zhang J, Wu J, Liu X, Xu J, Zhang J, et al. Disrupted neural variability during propofol-induced sedation and unconsciousness. Hum Brain Mapp. 2018;39(11):4533–4544. doi:10.1002/hbm.24304.

52. Husserl E. Inner time consciousness. Amsterdam:Nijhaus Publisher;1921.

53. Jack AI, Roepstorff A. Introspection and cognitive brain mapping: from stimulus-response to script-report. Trends Cogn Sci. 2002;6(8):333–339.

54. James W. The Principles of Psychology: Vol. 1. London: Macmillan;1890.

55. James W. The Principles of Psychology: Vol. 2. London: Macmillan;1890.

56. Kanai R, Tsuchiya N. Qualia. Curr Biol. 2012;22(10):R392–6. doi: 10.1016/j.cub.2012.03.033.

57. Kanwisher N, Yovel G. The fusiform face area: a cortical region specialized for the perception of faces. Philos Trans R Soc Lond B Biol Sci. 2006;361(1476):2109–28. doi: 10.1098/rstb.2006.1934

58. Kant I. Critique of Pure Reason. Guyer P, Wood AW, editors. Cambridge:Cambridge University Press:1781/1998.

59. Koch C. The Quest for Consciousness. Oxford:Oxford University Press; 2004.

60. Koch C, Massimini M, Boly M, Tononi G. Neural correlates of consciousness: progress and problems. Rev. Neurosci. 2016;17(5):307–21. doi: 10.1038/nrn.2016.22.

61. Koch C. What Is Consciousness? Nature. 2018;557(7704):S8–S12. doi: 10.1038/d41586-018-05097-x.

62. Lamme VA, Roelfsema PR. The distinct modes of vision offered by feedforward and recurrent processing. Trends Neurosci. 2000;23(11):571–9.

63. Lau H, Rosenthal D. Empirical support for higher-order theories of conscious awareness. Trends Cogn Sci. 2011;15(8):365–73. doi: 10.1016/j.tics.2011.05.009.

64. Levine J. Materialism and qualia: the explanatory gap. Pac Philos Q, 1983;64(4):354–361. doi: 10.1111/j.1468-0114.1983.tb00207.x

65. Linkenkaer-Hansen K, Nikouline VV, Palva JM, Ilmoniemi RJ. Long-range temporal correlations and scaling behavior in human brain oscillations. J Neurosci. 2001;21(4):1370–7. doi: 10.1523/JNEUROSCI.21-04-01370.2001

66. Liu CH, Ma X, Song LP, Fan J, Wang WD, Lv XY, et al. Abnormal spontaneous neural activity in the anterior insular and anterior cingulate cortices in anxious depression. Behav Brain Res. 2015;281:339–47. doi: 10.1016/j.bbr.2014.11.047.

67. Manning JR, Jacobs J, Fried I, Kahana MJ. Broadband shifts in local field potential power spectra are correlated with single-neuron spiking in humans. J Neurosci. 2009;29(43):13613–20. doi: 10.1523/JNEUROSCI.2041-09.2009.

68. Murray JD, Bernacchia A, Freedman DJ, Romo R, Wallis JD, Cai X, et al. A hierarchy of intrinsic timescales across primate cortex. Nat Neurosci. 2014;17(12):1661–3. doi: 10.1038/nn.3862.

69. Nagel T. What is it like to be a bat? Philos Rev. 1974;83(4):435–50. doi:10.2307/2183914

70. Northoff G. What the brain’s intrinsic activity can tell us about consciousness? A tri-dimensional view. Neurosci Biobehav Rev. 2013;37(4):726–38. doi:10.1016/j.neubiorev.2012.12.004

71. Northoff G. Unlocking the brain: Volume I: Coding. Oxford: Oxford University Press; 2014a.

72. Northoff G. Unlocking the brain: Volume II: Consciousness. Oxford: Oxford University Press; 2014b.

73. Northoff G. Neuro-philosophy and the healthy mind: Learning from the unwell brain. New York: Norton Publisher; 2016.

74. Northoff G. “Paradox of slow frequencies” – Are slow frequencies in upper cortical layers a neural predisposition of the level/state of consciousness (NPC)? Conscious Cogn. 2017;54:20–35. doi: 10.1016/j.concog.2017.03.006.

75. Northoff G. The spontaneous brain. From mind-body problem to world-brain problem. Cambridge: MIT Press; 2018.

76. Northoff G. The anxious brain and its heart – Temporal brain-heart de-synchronization in anxiety disorders. J Affective Disorders. Forthcoming 2019.

77. Northoff G, Huang Z. How do the brain’s time and space mediate consciousness and its different dimensions? Temporo-spatial theory of consciousness (TTC). Neurosci Biobehav Rev. 2017;80:630–645. doi: 10.1016/j.neubiorev.2017.07.013.

78. Oizumi M, Albantakis L, Tononi G. From the phenomenology to the mechanisms of consciousness: Integrated Information Theory 3.0. PLoS Comput Biol. 2014;10(5):e1003588. doi: 10.1371/journal.pcbi.1003588.

79. Oizumi M, Tsuchiya N, Amari SI. Unified framework for information integration based on information geometry. Proc Natl Acad Sci USA. 2016;113(51):14817–14822. doi: 10.1073/pnas.1603583113.

80. Oizumi M, Amari S, Yanagawa T, Fujii N, Tsuchiya N. Measuring integrated information from the decoding perspective. PLoS Comput Biol. 2016;12(1):e1004654. doi: 10.1371/journal.pcbi.1004654.

81. Palva JM, Zhigalov A, Hirvonen J, Korhonen O, Linkenkaer-Hansen K, Palva S. Neuronal long-range temporal correlations and avalanche dynamics are correlated with behavioral scaling laws. Proc Natl Acad Sci USA. 2013;110(9):3585–90. doi: 10.1073/pnas.1216855110.

82. Phillips S, Wilson WH. Categorial compositionality: a category theory explanation for the systematicity of human cognition. PLoS Comput Biol. 2010;6(7):e1000858. doi: 10.1371/journal.pcbi.1000858.

83. Phillips S, Wilson WH. Systematicity and a categorical theory of cognitive architecture: Universal construction in context. Front Psychol. 2016;7:1139. doi: 10.3389/fpsyg.2016.01139.

84. Ponce-Alvarez A, He BJ, Hagmann P, Deco G. Task-driven activity reduces the cortical activity space of the brain: Experiment and whole-brain modeling. PLoS Comput. Biol. 2015;11(8): e1004445. doi:10.1371/journal.pcbi.1004445

85. Rangarajan V, Hermes D, Foster BL, Weiner KS, Jacques C, Grill-Spector K, et al. Electrical stimulation of the left and right human fusiform gyrus causes different effects in conscious face perception. J Neurosci. 2014;34(38):12828–36. doi: 10.1523/JNEUROSCI.0527-14.2014.

86. Rees G, Friston K, Koch C. A direct quantitative relationship between the functional properties of human and macaque V5. Nat Neurosci. 2000;3(7):716–23. doi: 10.1038/76673

87. Rosenthal DM. Metacognition and higher-order thoughts. Conscious Cogn. 2000;9(2 Pt 1):231–42. doi: 10.1006/ccog.2000.0441

88. Sadaghiani S, Hesselmann G, Friston KJ, Kleinschmidt A. The relation of ongoing brain activity, evoked neural responses, and cognition. Front Syst Neurosci. 2010;4:20. doi: 10.3389/fnsys.2010.00020.

89. Sadaghiani S, Hesselmann G, Kleinschmidt A. Distributed and antagonistic contributions of ongoing activity fluctuations to auditory stimulus detection. J Neurosci. 2009;29(42):13410–7. doi: 10.1523/JNEUROSCI.2592-09.2009.

90. Sadaghiani S, Poline JB, Kleinschmidt A, D’Esposito M. Ongoing dynamics in large-scale functional connectivity predict perception. Proc Natl Acad Sci USA. 2015;112(27):8463–8. doi: 10.1073/pnas.1420687112.

91. Scalabrini A, Ebisch SJH, Huang Z, Di Plinio S, Perrucci MG, Romani GL, et al. Spontaneous Brain Activity Predicts Task-Evoked Activity During Animate Versus Inanimate Touch. Cereb Cortex. 2019. doi: 10.1093/cercor/bhy340.

92. Scalabrini A, Huang Z, Mucci C, Perrucci MG, Ferretti A, Fossati A, et al. How spontaneous brain activity and narcissistic features shape social interaction. Sci Rep. 2017;7(1):9986. doi: 10.1038/s41598-017-10389-9.

93. Schurger A, Sarigiannidis I, Naccache L, Sitt JD, Dehaene S. Cortical activity is more stable when sensory stimuli are consciously perceived. Proc Natl Acad Sci USA. 2015;112(16):E2083–92. doi: 10.1073/pnas.1418730112.

94. Searle JR. Mind: A Brief Introduction. Oxford: Oxford University Press; 2004.

95. Seth AK, Barrett AB. Neural theories need to account for, not discount, introspection and behavior. Cogn Neurosci. 2010;1(3):227–8. doi: 10.1080/17588928.2010.496533.

96. Sylvester CM, Shulman GL, Jack AI, Corbetta M. Anticipatory and stimulus-evoked blood oxygenation level-dependent modulations related to spatial attention reflect a common additive signal. J Neurosci. 2009;29(34):10671–82. doi: 10.1523/JNEUROSCI.1141-09.2009.

97. Tagliazucchi E, Chialvo DR, Siniatchkin M, Amico E, Brichant JF, Bonhomme V, et al. Large-scale signatures of unconsciousness are consistent with a departure from critical dynamics. J R Soc Interface 2016;13(114): 20151027. doi: 10.1098/rsif.2015.1027.

98. Tagliazucchi E, von Wegner F, Morzelewski A, Brodbeck V, Jahnke K, Laufs H. Breakdown of long-range temporal dependence in default mode and attention networks during deep sleep. Proc Natl Acad Sci USA. 2013;110:15419–24. doi:10.1073/pnas.1312848110

99. Tegmark M. Improved measures of integrated information. PLoS Comput Biol. 2016;12(11):e1005123. doi: 10.1371/journal.pcbi.1005123.

100. Tong F, Nakayama K, Vaughan JT, Kanwisher N. Binocular rivalry and visual awareness in human extrastriate cortex. Neuron. 1998;21(4):753–9.

101. Tononi G. An information integration theory of consciousness. BMC Neurosci. 2004;5:42. doi: 10.1186/1471-2202-5-42.

102. Tononi G, Boly M, Massimini M, Koch C. Integrated information theory: from consciousness to its physical substrate. Nat Rev Neurosci. 2016;17:450–61. doi: 10.1038/nrn.2016.44

103. Tononi G, Koch C. Consciousness: here, there and everywhere? Phil Trans R Soc B. 2015;370(1668). pii: 20140167. doi: 10.1098/rstb.2014.0167.

104. Tononi G, Edelman GM. Consciousness and complexity. Science. 1998;282(5395):1846–51.

105. Tsuchiya N, Taguchi S, Saigo H. Using category theory to assess the relationship between consciousness and integrated information theory. Neurosci Res. 2016;107:1–7. doi: 10.1016/j.neures.2015.12.007.

106. Vaadia E, Haalman I, Abeles M, Bergman H, Prut Y, Slovin H, et al. Dynamics of neuronal interactions in monkey cortex in relation to behavioural events. Nature. 1995;373(6514):515–8. doi: 10.1038/373515a0

107. Wolff A, Di Giovanni DA, Gómez-Pilar J, Nakao T, Huang Z, Longtin A, et al. The temporal signature of self: Temporal measures of resting-state EEG predict self-consciousness. Hum Brain Mapp. 2019a;40(3):789–803. doi:10.1002/hbm.24412.

108. Wolff A, Gómez-Pilar J, Nakao T, Northoff G. Interindividual neural difference in moral decision-making are mediated by alpha power and delta/theta phase coherence. Sci Rep. Forthcoming 2019b.

109. Zhang J, Magioncalda P, Huang Z, Tan Z, Hu X, Hu Z, et al. Altered global signal topography and its different regional localization in motor cortex and hippocampus in mania and depression. Schizophr Bull. 2018. doi:10.1093/schbul/sby138.

110. Zhou H, Melloni L, Poeppel D, Ding N. Interpretations of Frequency Domain Analyses of Neural Entrainment: Periodicity, Fundamental Frequency, and Harmonics. Front Hum Neurosci. 2016;10:274. doi: 10.3389/fnhum.2016.00274.

